# Circulating tumor cell heterogeneity and MMP9 expression in clinical correlation with breast cancer

**DOI:** 10.1101/2025.06.30.662312

**Authors:** Man Yi, Cuibin Sun, Minxing Liang, Jiahui Liu, Jiaxin Wei, Jinping Long, Haiqi Zhao, Shengtao Liu, Manyuan Tang, Zhongcheng Mo, Xin Li, Yue Yin

## Abstract

**Objectives:** This study aimed to investigate the association between the heterogeneity of circulating tumor cell (CTC) subtypes and the expression patterns of matrix metalloproteinase 9 (MMP9) with clinicopathological parameters in breast cancer patients. The distribution of CTC subtypes, MMP9 expression levels, and their influence on tumor progression.

**Methods:** A total of 303 breast cancer patients were enrolled in this study. The association between CTCs subtypes in peripheral blood, MMP9 expression and clinical or pathological data of patients were analyzed.

**Results:** The number of mesenchymal CTCs increased significantly with the number of lymph node metastases (*P* < 0.05), and high MMP9 expression was positively correlated with lymph node involvement (*P* < 0.05). The proportion of intermediate to high MMP9 expression was significantly higher in mixed CTCs compared to epithelial and mesenchymal CTCs (*P* < 0.05). High MMP9 expression in tumor tissues was associated with increased CTCs production and induction of epithelial-mesenchymal transition (EMT) (*P* < 0.05), as well as an elevated number of MMP9-positive CTCs (*P* < 0.05).

**Conclusions:** These findings suggest that MMP9 is positively associated with CTCs production and EMT process, and MMP9 may contribute to tumor metastasis by promoting CTC shedding and EMT by degrading the extracellular matrix.

We recommend compliance with the Mesh guidelines: https://www.ncbi.nlm.nih.gov/mesh/

## Introduction

Breast cancer has the highest incidence rate among women worldwide, and its incidence rate continues to climb, especially among younger age groups [1–4]. Breast cancer has the potential to metastasize via both lymphatic and hematogenous routes even at early stages. The most common sites of metastasis include the bones, lungs, liver, and brain [5–7]. CTCs shed from primary tumors or metastatic sites, can enter the peripheral circulation and disseminate to colonize distant organs. CTCs thus represent the dynamic biomarkers of tumor invasion and metastatic progression [8,9]. EMT is a mechanism that contributes to tumor development and metastasis by inhibiting apoptosis, evading immune elimination and promoting basement breaching and migration of CTCs [10–12].

MMPs constitute a family of zinc-dependent endopeptidases that degrade extracellular matrix constituents. These enzymes participate in physiological processes while contributing to pathological conditions such as arthritis and oncogenesis. MMPs facilitate tumor invasion and metastasis through three principal mechanisms: extracellular matrix degradation, activation of oncogenic signaling pathways, and induction of EMT. Among this family, MMP9 increases genetic susceptibility to type 2 diabetes through its gene polymorphisms, while its overexpression in hyperglycemic microenvironments exacerbates complications by degrading extracellular matrix and inhibiting angiogenesis; however, this characteristic can be exploited for developing MMP-9-responsive smart hydrogels to facilitate treatment [13,14], MMP9 also plays a critical role in metastatic progression and serves as a key clinical biomarker for assessing breast cancer prognosis [15–18]. However, the mechanistic relationship linking MMP9 and CTCs in clinical samples is not fully elucidated.

This study characterized subtypes of peripheral blood CTCs, total CTC counts, and MMP9 expression in breast cancer patients. Our study systematically assessed the correlations between CTC heterogeneity and clinicopathological characteristics, associations of MMP9 expression with clinicopathological parameters, and interrelationships among CTC subtypes, total CTC counts, and MMP9 expression. These findings establish a theoretical framework for developing metastasis risk stratification models and optimizing personalized targeted therapies.

## Materials and methods

### Patients

A total of 303 breast cancer patients were included in this study (Table 1). Blood or tissue samples were collected for CTC analyses or immunohistochemical staining (Table 1). All procedures involving pathological samples and patient record retrospect were according to the guideline of Ethics Review Board of Nanfang hospital. Pathological data, such as patients age, sex, tumor diameter, lymph node metastasis, Ki-67, and HER-2 status were recorded in the study. Clinical staging was determined according to the 8th edition of the AJCC breast cancer staging system. This study was approved by the Ethics Committee of Nanfang Hospital and written consent forms were obtained from patients.

**Table 1:**
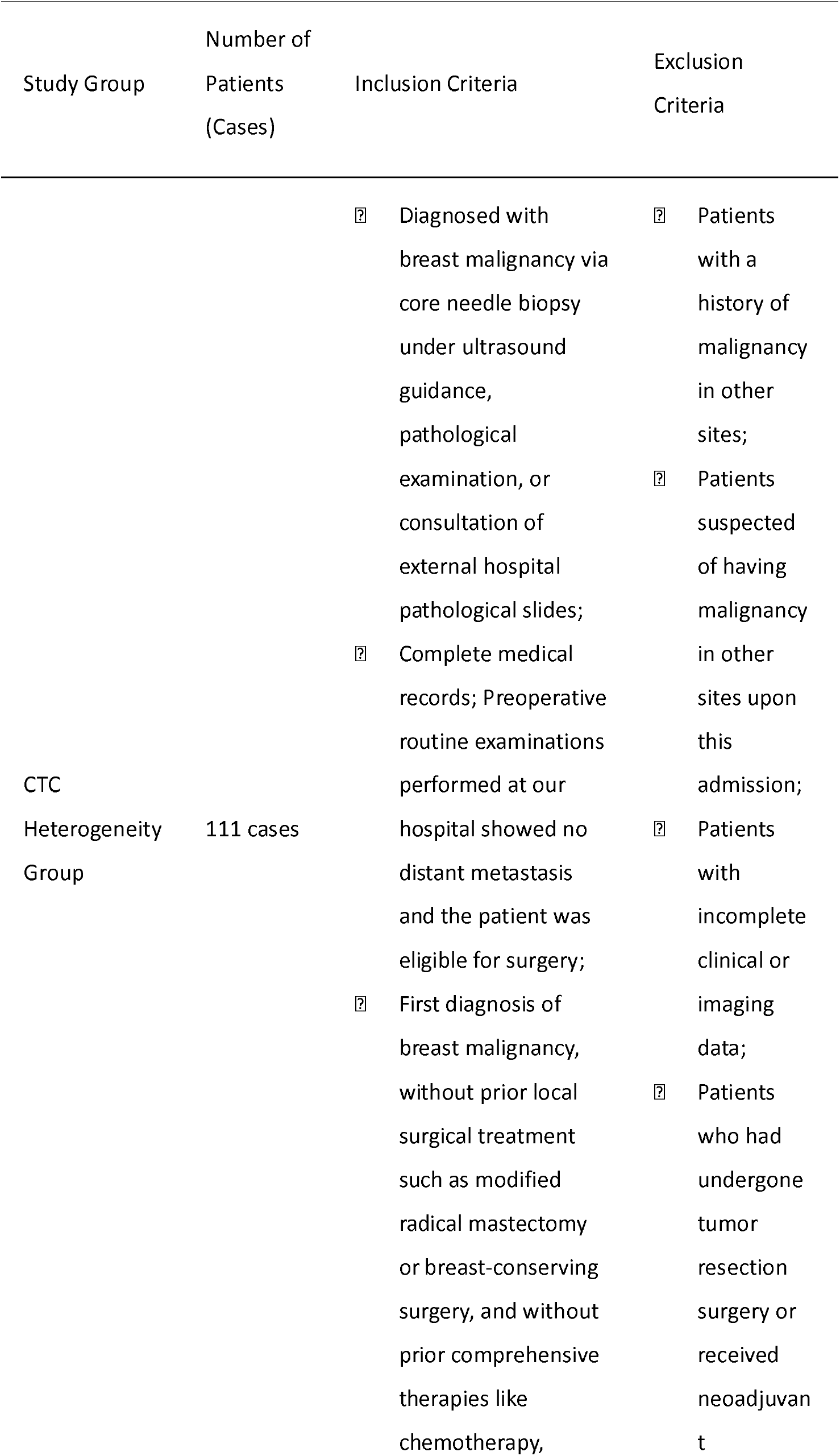
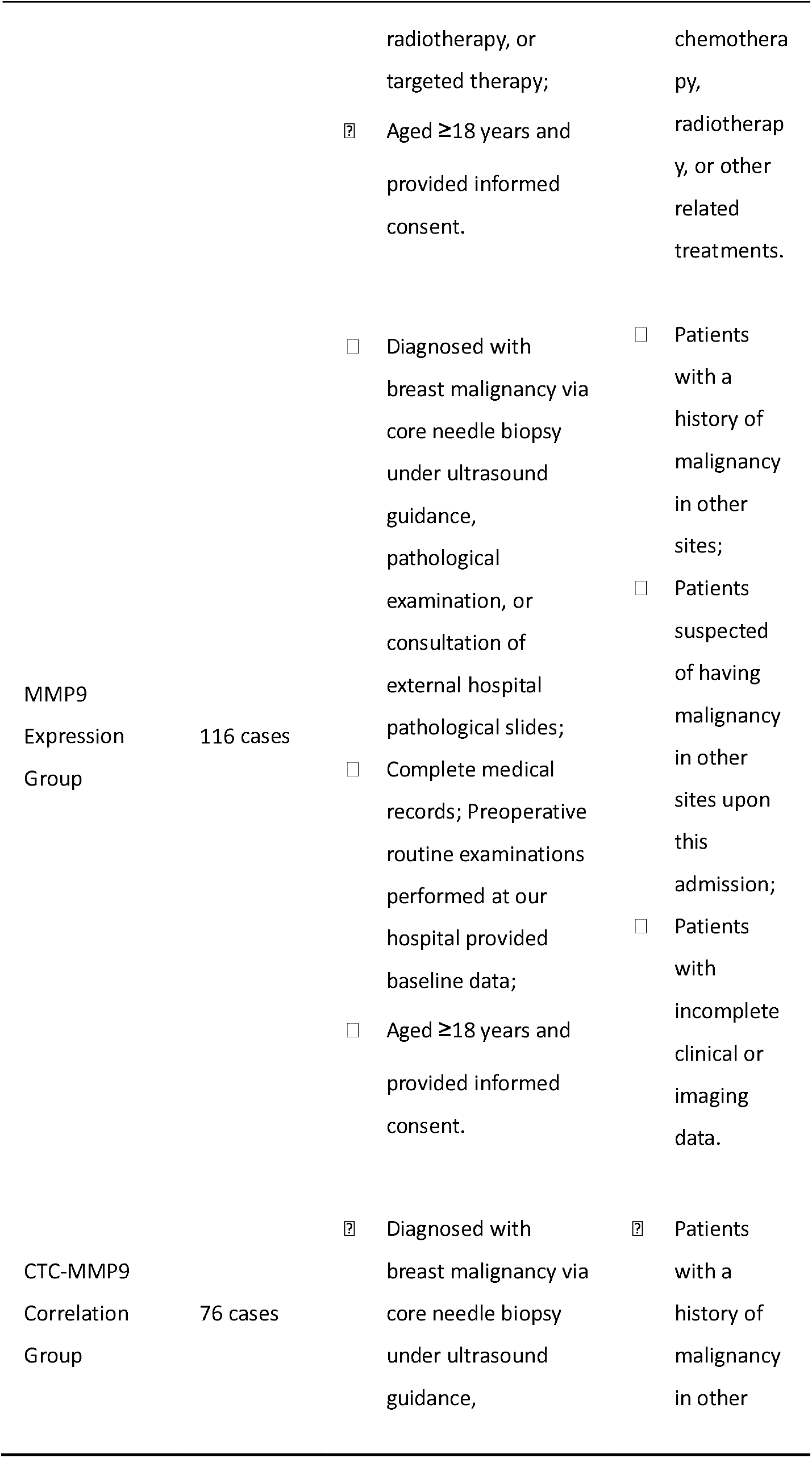
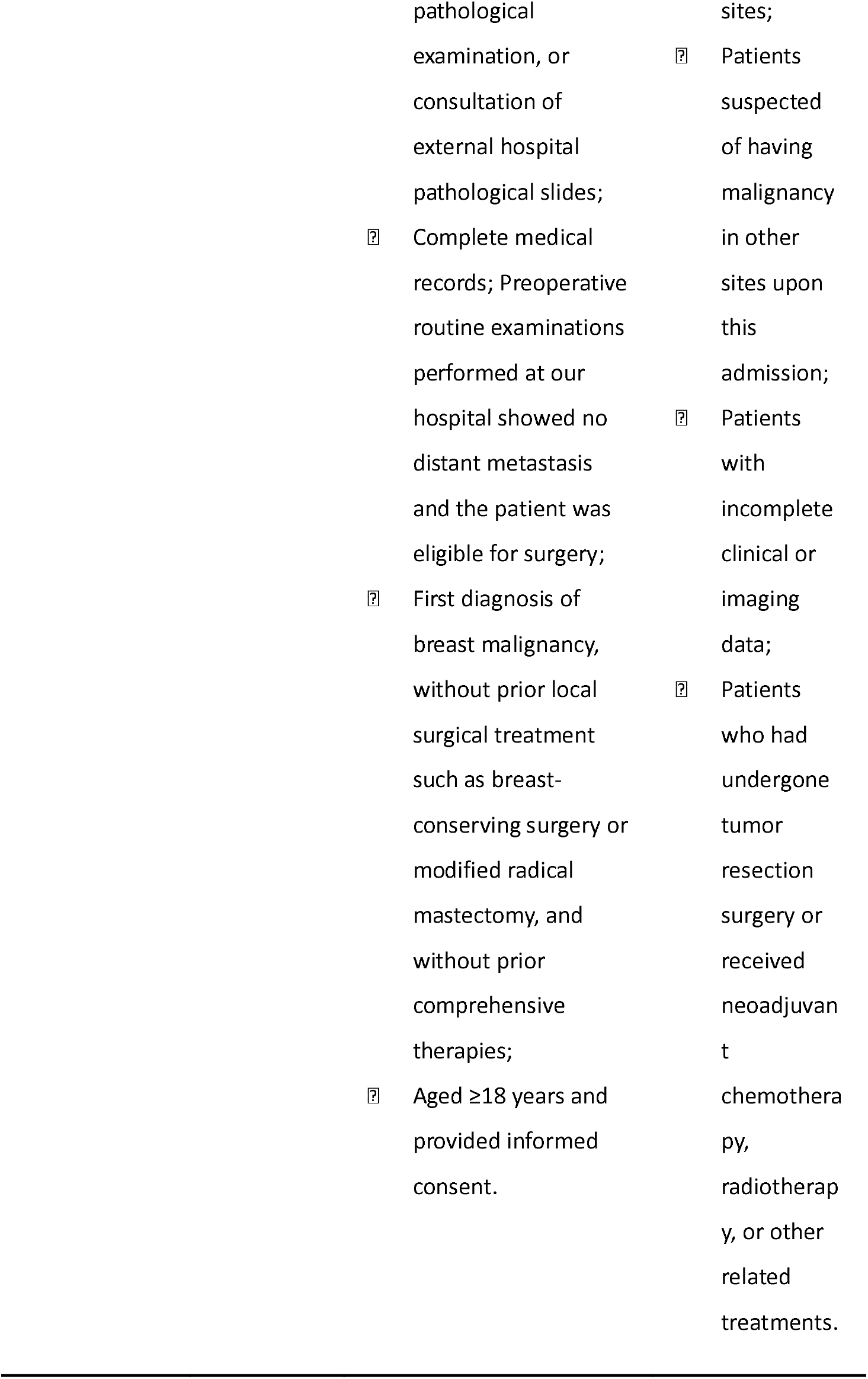
Patients Characteristic.

### CTCs detection

Peripheral venous blood was collected early in the morning before surgery, and CTCs were detected by CanPatrol™ second-generation CTC detection technology [19]. Briefly, Canpatrol™ CTC detection technology consists of two parts: CTC isolation and CTC classification and identification. Red blood cells in the peripheral blood were first lysed, and then CTCs were separated and enriched using nanotechnology based on the size differences between CTCs and white blood cells. In the CTC classification and identification part, a novel multiple mRNA in situ analysis method was used to specifically locate nucleic acids of enriched CTCs, achieving the purpose of classifying and identifying CTCs. This method used multiple RNA probes targeting CTCs simultaneously, and after hybridization with the target genes, the fluorescence signal amplification system could increase detection sensitivity to single-copy mRNA. C Epithelial markers (EpCAM, CK8/18/19) were labeled with Alexa Fluor 594 (red), while mesenchymal markers (Twist and Vimentin) were labeled with Alexa Fluor 488 (green). Leukocytes were identified by CD45 (Alexa Fluor 750, white) (Table S1). CTC subtypes were categorized based on fluorescence: epithelial (red+), mesenchymal (green+), and mixed (red+/green+). Additionally, MMP9 expression was detected using an Alexa Fluor 647 (purple) labeled probe. All CTC subtypes were quantified independently.

### Immunohistochemical (IHC) staining and fluorescence in situ hybridization (FISH)

Breast cancer tissue specimens were fixed in 10% formaldehyde and embedded in paraffin. Sections (4 μm thick) were deparaffinized, rehydrated, and subjected to heat-induced antigen retrieval in citrate buffer (pH 6.0). The sections were incubated overnight at 4°C with primary monoclonal antibodies against Ki-67 (cat. TA500265), HER-2 (cat. ZM-0065), and MMP9 (cat. TA336901) (all from Beijing Zhongshan Jinqiao Biotechnology Co., Ltd.). Subsequently, sections were incubated with a biotin-labeled Rabbit Anti-Mouse Secondary Antibody (cat. DS-0004, Beijing Zhongshan Jinqiao Biotechnology Co., Ltd.) at room temperature for 45 minutes. Visualization was performed using DAB Staining Reagent (cat. P0203, Beyotime Biotech Co., Ltd.), followed by hematoxylin counterstaining. Negative controls were prepared by replacing the primary antibody with PBS.

The slides were analyzed in a double-blind manner by two independent pathologists. Scores were assigned based on the intensity of positive staining: 1 for weak positivity (+), 2 for moderate positivity (++), and 3 for strong positivity (+++). The formula for scoring was calculated as follows: (+) %×1 + (++) %×2 + (+++) %×3. A total score of less than 1.0 was classified as (+), 1.0 to 1.5 as (++), and greater than 1.5 as (+++).

To determine HER2 status in breast cancer, 2013 the American Society of Clinical Oncology (ASCO) and the College of American Pathologists (CAP) guidelines were followed [20]. HER-2 (+) is negative and HER-2 (+++) is positive. When HER-2 expression cannot be determined by IHC (HER-2++), additional FISH should be done via HER-2/CEP-7 ratio. HER-2/CEP17 ratio ≥ 2.0 or mean HER-2 copy number/cell ≥ 6.0 was determined as HER-2 positive; HER-2/CEP17 ratio < 2.0 and mean HER-2 copy number/cell < 4.0 was determined as HER-2 negative. HER-2/CEP17 ratio < 2.0, mean HER-2 copy number/cell < 6.0, and ≥ 4.0 could not be determined as HER-2 results, combined with IHC results or re-selected other tissue blocks for examination; HER-2/CEP17 ratio ≥ 2.0 and mean HER-2 copy number/cell < 4.0, was controversial and required further interpretation based on the immunohistochemistry results, along with communication with the patient (representative FISH image of HER-2 positive is in Fig. S1f).

Ten consecutive fields of view were observed under a 400x microscope to derive the respective proportions of MMP9 no expression, low expression, medium expression, and high expression, which were multiplied by their weights. The sums were (+) for less than 1.0, (++) for between 1.0 and 1.5, and (+++) for greater than 1.5.

### Statistical software and methods

Statistical analyses were performed using IBM SPSS Statistics for Windows, version 22.0 (IBM Corp., Armonk, N.Y., USA) software. Comparisons between groups were conducted using the independent samples t-test, ANOVA, Welch’s ANOVA (for groups with unequal variances), or nonparametric tests. For Welch’s ANOVA, when significant differences among groups were detected (*P* < 0.05), post hoc comparisons were performed using the Games-Howell test to identify specific group differences. For ANOVA with equal variances assumed, the Tukey’s honestly significant difference (HSD) test was used for post hoc comparisons. Categorical variables were presented as frequencies (percentages), and the chi-square test or Fisher’s exact test was applied. *P* < 0.05 was considered statistically significant.

## Results

### Correlation of CTCs heterogeneity with clinical features

A total of 111 patients with invasive breast cancer were enrolled in this phase of the study (Table 1), with a mean age of 49.82 ± 9.63 years, ranging from 28 to 80 years. The clinical stages were as follows: 34 patients (30.6%) were in stage IA, 20 patients (18%) were in stage IB, 32 patients (28.8%) were in stage IIA, 11 patients (9.9%) were in stage IIB, 5 patients (4.5%) were in stage IIIA, 7 patients (6.3%) were in stage IIIB, and 2 patients (1.8%) were in stage IIIC. Among these patients, 110 were female and one was male.

CTCs were identified in 99 patients (89.19%), with a total of 593 CTCs detected, including 144 epithelial CTCs (24.28%), 319 mixed CTCs (53.79%), and 130 mesenchymal CTCs (21.92%). The mean total number of CTCs was 5.31±5.89, with 1.2±1.66 epithelial CTCs, 2.86±3.79 mixed CTCs, and 1.16±2.20 mesenchymal CTCs (Fig. 1A).

**Figure 1:**
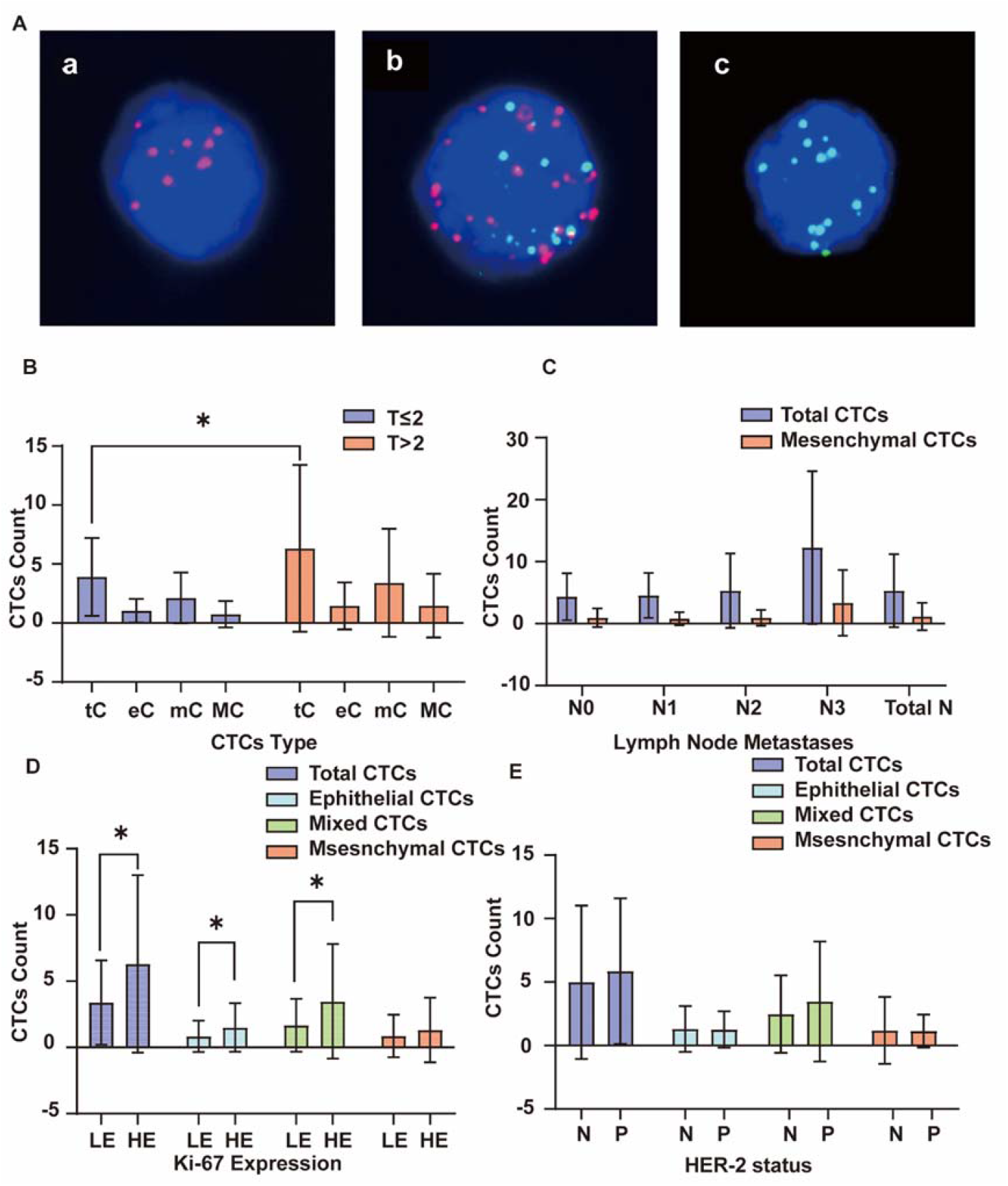
Correlation of circulating tumor cell (CTC) heterogeneity with clinical features. (A) a: epithelial CTC (red fluorescence signal); b: mixed CTC (mixed red and green fluorescence signals); c: mesenchymal CTC (green fluorescence signal). (B) The distribution of total CTCs (tC) epithelial CTCs (eC) mixed CTCs (mC) and mesenchymal CTCs (MC) in patients with tumor diameter ≤ 2 cm and > 2 cm. Data show significantly lower total CTCs in the ≤ 2 cm group (CTCs were quantified as counts/5ml. \**P* < 0.05, independent samples t-test). Error bars represent SD values. (C) Distribution of total CTCs (tc) and CTC subtypes across lymph node metastasis groups. N0: No regional lymph node metastasis. N1: Metastasis in ipsilateral peribronchial and/or ipsilateral hilar lymph nodes, and intrapulmonary lymph node metastasis. N2: Metastasis in ipsilateral mediastinal and/or subcarinal lymph nodes. N3: Metastasis in contralateral mediastinal, contralateral hilar, ipsilateral or contralateral scalene, and supraclavicular lymph nodes. The Kruskal-Wallis test revealed significant differences in total and mesenchymal CTC distributions among groups (*P* < 0.05). Error bars represent SD values. (D) Comparison of total CTCs and CTC subtypes between patients with low (LE, < 20%) and high (HE, ≥ 20%) Ki-67 expression. Total CTCs, epithelial CTCs, and mixed CTCs were significantly higher in the HE group (all * *P* < 0.05, independent samples t-test). Error bars represent SD values. (E) Comparison of total CTCs and CTC subtypes between HER-2 negative (N) and positive (P) groups. No significant differences were observed (*P* > 0.05, independent samples t-test). Error bars indicate SD. Among the patients, one case with HER-2 (++) did not undergo FISH testing; two patients with intraductal carcinoma were not suitable for HER-2 testing. Error bars represent SD values.

To explore potential clinical correlates, we assessed the relationship between CTC counts and clinical stage. There were significant differences in the distribution of the total number of CTCs and mixed CTCs at different clinical stages (*P* < 0.05). Although the mean values of epithelial and mesenchymal CTCs were higher in stages II and III than those in stage I, the differences between the different clinical stages were not statistically significant (*P* > 0.05; Table 2).

**Table 2:** Comparison of CTCs distribution in different clinical stages.

| Clinical Stage | Stage I (n=54) | Stage II/III (n=57) | *t* | *P* |
| --- | --- | --- | --- | --- |
| Total CTCs | 3.93±3.34 | 6.61±7.35 | 7.7 | 0.014* |
| Epithelial CTCs | 1.07±1.20 | 1.47±1.99 | 6.02 | 0.201 |
| Mixed CTCs | 2.04±2.16 | 3.65±4.74 | 10.62 | 0.023* |
| Mesenchymal CTCs | 0.81±1.08 | 1.49±2.85 | 4.65 | 0.1 |

CTCs units: cells/5mL, expressed as mean ± standard deviation; \* indicates $P < 0.05$ , indicating a statistically significant difference.

Total number of CTCs in the group with a tumor diameter > 2 cm was statistically higher than that in the group with a tumor diameter ≤ 2 cm (*P* < 0.05). CTCs subtypes were not significantly differentially distributed between the two groups (all *P* > 0.05; Fig. 1B). Subsequently, the potential link with lymph node metastasis burden was examined. A total of 111 patients were categorized into the N0 group (64, 57.7%), N1 group (20, 18.0%), N2 group (16, 14.4%), and the N3 group (11, 9.9%) according to lymph node metastasis (Table 1). The Kruskal-Wallis test revealed significant differences in the distribution of total CTCs and mesenchymal CTCs among different lymph node metastasis groups (all *P* < 0.05), with both increasing as lymph node metastasis progressed. However, no statistically significant differences were observed in epithelial or mixed CTCs (both *P* > 0.05; Table 3).

**Table 3:** Comparison of CTCs distribution among different lymph node metastasis groups.

| Regional Lymph Nodes | N0<br>(n=64) | N1<br>(n=20) | N2<br>(n=16) | N3<br>(n=11) | Z | P |
| --- | --- | --- | --- | --- | --- | --- |
| Total CTCs | 3(0,14) | 4(0,14) | 4(0,25) | 6(3,43) | 7.983 | 0.046* |
| Epithelial CTCs | 1(0,5) | 0.5(0,5) | 0.5(0,3) | 1(0,11) | 4.278 | 0.233 |
| Mixed CTCs | 1(0,9) | 2(0,13) | 2(0,22) | 3(0,21) | 4.887 | 0.18 |
| Mesenchymal CTCs | 0(0,9) | 0(0,3) | 0(0,4) | 2(0,19) | 9.371 | 0.025* |

CTCs units: cells/5mL, expressed as median (minimum, maximum); \* indicates $P < 0.05$ , indicating a statistically significant difference between groups. N1: Metastasis in 1-3 ipsilateral axillary lymph nodes; N2: Metastasis in 4-9 ipsilateral axillary lymph nodes; N3: Metastasis in ≥10 ipsilateral axillary lymph nodes.

Among the 111 patients (Table 1), 38 patients (34.2%) were in the low-expression group, and 73 patients (65.8%) were in the high-expression group based on Ki-67 expression levels (20%) (representative Ki-67 IHC images in Fig. S1a-b). Significant differences were observed in the distribution of the total number of CTCs, as well as in the distribution of epithelial and mixed CTCs between the two groups (*P* < 0.05; Fig. 1D).

In contrast to Ki-67 results, HER-2 status did not appear to strongly correlate with CTC profiles. There were 67 patients (60.91%) who were HER-2 negative and 43 HER-2 positive patients (39.09%) (Table 1) (representative HER-2 IHC/FISH images in Fig. S1c-f). There were no significantly significant differences in the total number of CTCs or the distribution of any CTC subtypes between the HER-2 positive and negative groups (all *P* > 0.05; Fig. 1E). Finally, we investigated whether CTC patterns differed across intrinsic molecular subtypes of breast cancer. Patients were grouped according to the expression of HER-2, ER, PR, and Ki-67 status, with 30 cases of luminal A type (27.03%), 28 cases of luminal B HER-2 (-) type (25.22%), 26 cases of luminal B HER-2 (+) type (23.42%), HER-2 positive type was found in 17 cases (15.32%) and triple-negative type was found in 9 cases (8.11%). Because one patient showed (++) on HER-2 examination, FISH examination was not performed. This patient was not included in the subsequent statistical analyses. Kruskal-Wallis test showed that the total number of CTCs and the distribution of each subtype did not differ statistically among the different molecular typing groups (all *P* > 0.05; Table 4).

**Table 4:** Comparison of CTCs distribution among different molecular type groups.

| Molecular Type | Total CTCs | Epithelial CTCs | Mixed CTCs | Mesenchymal CTCs |
| --- | --- | --- | --- | --- |
| Luminal A | 3 (0,12 ) | 0 (0,5) | 1 (0,9) | 0 (0,9) |
| Luminal B HER-2 (-) Type | 4.5 (0,43) | 1 (0,11) | 2 (0,13) | 1 (0,19) |
| Luminal B HER-2 (+) Type | 3 (0,25) | 1 (0,5) | 1 (0,22) | 0.5 (0,4) |
| HER-2 Positive Type | 6 (1,25) | 1 (0,5) | 3 (0,21) | 2 (0,5) |
| Triple Negative Type | 4 (0,7) | 1 (0,4) | 2 (0,4) | 0 (0,1) |
| Z | 8.447 | 6.783 | 6.961 | 6.878 |
| P | 0.077 | 0.148 | 0.138 | 0.142 |

One patient whose HER-2 test showed (++), but did not undergo supplementary FISH testing, was therefore excluded from the statistical grouping during analysis.

### Correlation between MMP9 expression and clinical features

A total of 116 breast cancer patients were enrolled, with an age range from 30 to 80 years and a median age of 50.56±9.66 years (Table 1). Among these, 50 patients (43.10%) were lymph node-positive and 66 patients (56.90%) were lymph node-negative. The clinical stages were as follows: 2 patients (1.72%) were in Stage 0, 57 patients (49.14%) were in Stage I, 41 patients (35.34%) were in Stage II, 15 patients (12.93%) were in Stage III, and 1 patient (0.86%) was in Stage IV (Table 1).

Of the 116 patients (Table 1), 15 (12.9%) were weakly positive (+); 53 (45.7%) were moderately positive (++); and 48 (41.4%) were strongly positive (+++) (Fig. 2A).

**Figure 2:**
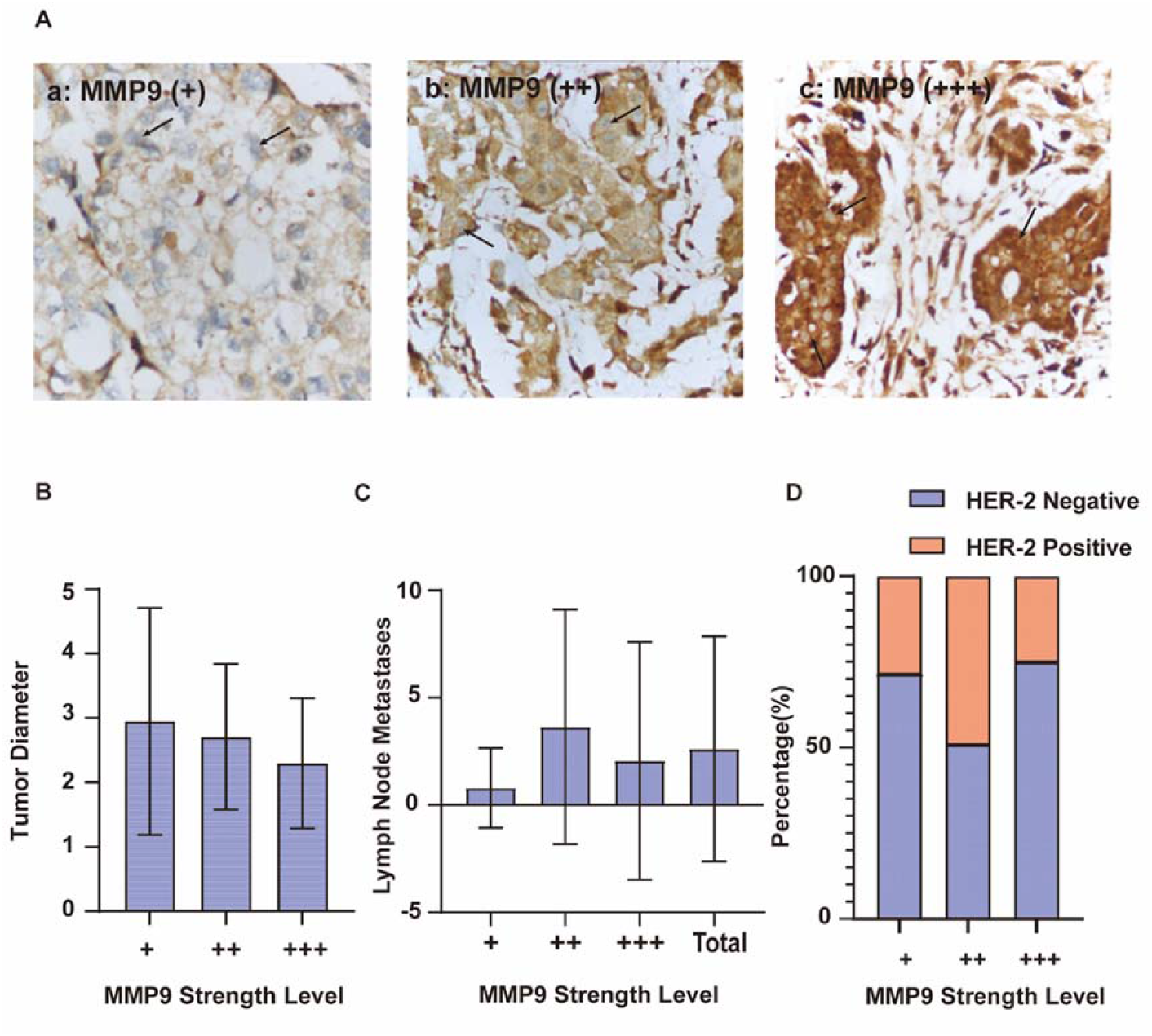
Correlation between matrix metalloproteinase-9 (MMP9) expression and clinical features. (A) Representative immunohistochemical staining of MMP9 expression levels: weak (+), moderate (++), or strong (+++). (B) The mean tumor diameter was significantly smaller in the high MMP9 expression group (+++) compared to the combined low/moderate expression group (+/++) (*P* = 0.039, independent samples t-test). Error bars represent SD values. (C) Lymph node metastasis counts across MMP9 expression groups (+), (++), (+++). Welch’s ANOVA (used due to unequal variance) showed a significant difference among groups (*P* < 0.05). Post hoc Games-Howell tests indicated fewer metastases in the (+++) group versus (+) group (*P* < 0.05). Error bars represent SD values. (D) A significant association was observed between MMP9 expression levels and HER-2 status (*P* = 0.037, Pearson’s χ^2^ test). One patient with HER-2 (++) did not undergo FISH testing, and two patients with DCIS were excluded from HER-2 assessment.

Based on these expression levels, patients were categorized into two groups for comparison: the high-expression group (+++), which included 48 cases (41.38%) with a mean tumor diameter of 2.30±1.01 cm, and the low- and medium-expression group (+) and (++), which included 68 cases (58.62%) with a mean tumor diameter of 2.76±1.28 cm. Importantly, mean tumor diameter in the high-expression group was significantly smaller than in the low- and medium-expression groups (*P* < 0.05; Fig. 2B).

Across the MMP9 expression groups (+), (++), (+++), the number of lymph node metastases was 0.8±1.859 (MMP9+), 3.64±5.463 (MMP9++), and 2.06±5.533 (MMP9+++). Variance homogeneity was assessed using Levene’s test (*F* = 3.266, *P* = 0.042), indicating unequal variances across groups; thus, Welch’s ANOVA (suitable for unequal variances) was performed. The Welch’s ANOVA revealed a statistically significant difference in lymph node metastases among the three groups (*F* = 5.156, P = 0.008). Post hoc comparisons using the Games-Howell test identified that the MMP9+++ group had significantly fewer lymph node metastases compared to the MMP9+ group (*P* < 0.05); the MMP9++ group also differed significantly from the MMP9+ group (*P* < 0.05).

Across the MMP9 expression groups (+), (++), (+++), the number of HER-2-negative patients consistently outnumbered that of HER-2-positive patients. Although the absolute number of negative patients increased with higher MMP9 expression levels, the proportion of HER-2-negative patients did not show a corresponding upward trend. Nevertheless, Statistical significance was observed among the groups (*P* < 0.05; Fig. 2D).

### Relationship between CTCs and MMP9 expression

A total of 76 patients with malignant breast tumors were included in the study (Table 1), with a mean age of 49.83±8.76 years, ranging from 30 to 69 years. Clinical stages were distributed as follows: Stage 0: 2 (2.63%), IA: 24 (31.58%), IB: 12 (15.79%), IIA: 26 (34.21%), IIB: 5 (6.58%), IIIA: 1 (1.32%), and IIIB: 6 (7.89%). Immunohistochemical examination of MMP9 in pathological sections revealed 19 cases (25%) of weak positivity (+), 45 cases (59%) of moderate positivity (++), and 12 cases (16%) of strong positivity (+++) in total (Table 1).

In both total CTCs and all CTC subtypes, CTCs counts based on MMP9 expression in CTCs (from negative to high) showed a progressive decrease (Fig. 3A, Fig. S2). We next analyzed CTC counts based on tumor tissue MMP9 status. Patients were divided into MMP9 (+) group (low-expression group) (19, 25.0%) and MMP9 (++) (+++) group (high-expression group) (57, 75.0%). The number of total CTCs and mixed CTCs was significantly higher in the high-expression group (both *P* < 0.05; Fig. 3B), and there was no significant difference between the epithelial and mesenchymal CTCs groups (both *P* > 0.05; Fig. 3B).

**Figure 3:**
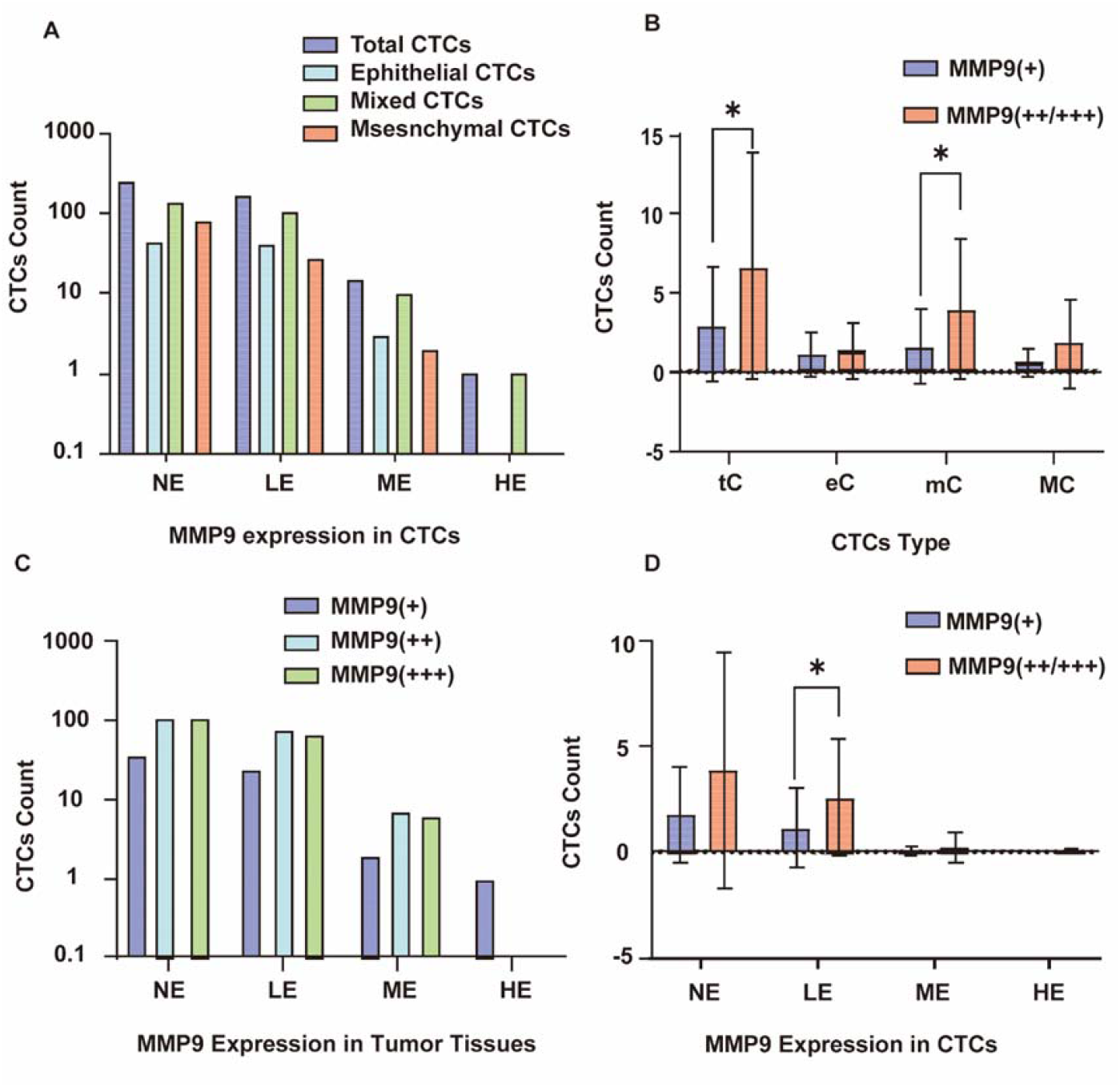
Relationship between CTCs and MMP9 expression. (A) Counts of each CTC subtype under different MMP9 expression levels. MMP9 expression levels were categorized as no expression (NE), low expression (LE), medium expression (ME), or high expression (HE). CTC counts were log10-transformed and plotted on a scale from 0.1 to 1000. (B) The distribution of total CTCs (tc), epithelial CTCs (eC), mixed CTCs (mC), and mesenchymal CTCs (MC) correlate with tumor MMP9 expression levels. Mixed CTCs and total CTCs were significantly higher in MMP9 (++/+++) than (+) groups (mixed CTCs: *t*=4.017, * *P* < 0.05, Welch’s t-test; total CTCs: * *P* < 0.05, standard t-test). Error bars represent SD values. (C) Analysis of MMP9 expression on CTCs (NE, LE, ME, HE) relative to tumor MMP9 expression levels. NE: no expression, LE: low expression, ME: medium expression, HE: high expression. (D) When CTCs showed low MMP9 expression, tumors with high MMP9 expression contained significantly more CTCs than those with low MMP9 expression (\**P* < 0.05, independent samples t-test). Error bars represent SD values.

After grouping patients according to the strength of MMP9 expression in tumor tissues, analysis revealed that the frequency of MMP9 expression in CTCs decreased sequentially from no expression to high expression. The proportion of MMP9 expression in tumor tissues that constituted different levels of MMP9 expression in CTCs, MMP9 (++) and (+++) accounted for a relatively high percentage of the total number of MMP9 (++) and (+++). However, the chi-square test showed that the difference between the groups was not statistically significant (Pearson χ^2^ = 6.306, *P* = 0.39; Fig. 3C). Based on the grouping of MMP9 expression levels on CTCs, tumor tissue MMP9 was divided into a low expression group (MMP9 (+) group) and high expression group (MMP9 (++) (+++) group). Independent samples t-tests showed that the number of CTCs in the tumor tissue MMP9 high expression group was statistically higher than in the low-expression group when CTCs were lowly expressed in MMP9 (*P* < 0.05; Fig. 3D).

## Discussion

The EMT phenotype of CTCs has been clinically validated as an important prognostic indicator for breast cancer outcomes [21–23]. Kalavska [24] employed RT-qPCR to quantify EMT-related transcripts in CTCs, revealing elevated MMP9 expression in aggressive breast cancer subtypes with potential clinical relevance. This study employed CanPatrol™ technology to achieve multidimensional phenotypic profiling of CTCs, overcoming limitations inherent to traditional CTC subtyping methodologies. This approach provides enhanced resolution of tumor heterogeneity and metastatic propensity. Furthermore, the simultaneous analysis of MMP9 expression in CTCs and tumor tissues revealed a correlation between CTC abundance and MMP9 expression.

Mesenchymal CTCs were prevalent in breast cancer patient groups and demonstrated significant correlation with metastatic progression. Pereira-Veiga [21] demonstrated that mesenchymal CTCs exhibit heightened invasive capacity but reduced proliferative potential. Ki-67, a key indicator of proliferative activity in breast cancer [25–27], has a central role in molecular typing, prognostic stratification, and treatment selection. In contrast, our analysis revealed significantly elevated CTC burden and increased proportions of mixed CTCs in patients with high Ki-67 expression, suggesting proliferative activity may potentiate CTC dissemination through enhanced tumor cell detachment or EMT induction. Detection of mixed and mesenchymal CTC phenotypes not only confirms active EMT in breast cancer, but also enables multimodal assessment of tumor progression, metastatic propensity, and proliferative activity through quantitative and phenotypic CTC dynamics. Elevated CTCs burden in peripheral blood has been documented in HER2-positive breast cancer patients [28]; however, this study found no significant association between HER-2 status or molecular subtypes and CTC distribution. This null association may be attributable to limited sample size or intrinsic heterogeneity in breast cancer phenotypes. Although this component of the investigation employed a cross-sectional design that captures only a momentary assessment of EMT status, it establishes a foundational framework for future longitudinal studies evaluating dynamic correlations between CTC profiles and clinical prognosis.

We also found that MMP9 expression intensity was positively correlated with the number of lymph node metastases. Furthermore, study has demonstrated that high MMP9 expression is associated with lymph node metastasis and poorer prognosis in breast cancer [29]. Paradoxically, tumors exhibiting strong MMP9 expression demonstrated significantly smaller diameters compared to moderately or weakly positive counterparts. This inverse correlation potentially reflects enhanced local invasive activity at tumor margins, constraining overall tumor expansion. Alternatively, limitations in immunohistochemical scoring system could contribute to this observation. The observed phenomenon suggests a dichotomous role for MMP9 in oncogenesis: in highly expressing tumors, MMP9 may facilitate hematogenous dissemination through activation of invasion-related signaling cascades, while tumors with attenuated MMP9 expression appear to favor primary tumor expansion, potentially through augmented proliferative signaling. This hypothesis is substantiated by clinical findings demonstrating attenuated proliferative activity in tumor cells with elevated MMP9 expression [24]. Consequently, we postulate that MMP9 may function as a “metastatic switch”, whose expression threshold governs the transition from localized tumor proliferation to systemic dissemination.

This investigation demonstrated that elevated MMP9 expression in both tumor tissues and CTCs facilitates CTC detachment from primary sites and potentiates EMT activation. Similarly, elevated MMP9 expression in hepatocellular carcinoma (HCC) also promotes EMT activation and facilitates tumor cell invasion through basement membrane degradation [30]. MMP9 deficiency dysregulates actin polymerization-related proteins, impairs cell migratory capacity, and attenuates TGF-β-induced EMT in lens epithelial cells [31]. It is well-established that EMT critically contributes to metastatic invasion in breast cancer [32–35]. Shen et al. further demonstrated that 27-hydroxycholesterol promotes EMT via MMP9 upregulation [36], establishing a mechanistic pathway that leads to breast cancer invasion. Therefore, MMP9 represents a critical molecular determinant modulating phenotypic reprogramming in both EMT and CTCs, with its expression levels demonstrating significant correlations with CTC subtyping, quantitative abundance, and metastatic efficiency.

This study has several limitations. Although 303 patients were enrolled, group analyses likely had reduced statistical power to detect subtle associations. Additionally, the geographic and ethnic homogeneity of the cohort may restrict the applicability of the results to populations with different genetic backgrounds or healthcare settings. Although significant correlations were observed between MMP9 expression and CTC characteristics, the underlying mechanisms remain speculative. Functional studies are needed to determine whether MMP9 directly drives EMT in CTCs or merely correlates with aggressive tumor phenotypes. Furthermore, the analysis did not account for intrinsic molecular subtypes of breast cancer. Since these subtypes exhibit distinct biological behaviors and metastatic potentials, the observed relationships between CTC heterogeneity and MMP9 expression could be influenced by subtype-specific mechanisms. Future studies incorporating subtype-stratified analyses are warranted to clarify these potential differences.

In summary, CTC heterogeneity and MMP9 overexpression demonstrate significant associations with clinicopathological characteristics in breast cancer. MMP9 may potentiate CTC metastatic competence through EMT induction while representing a critical regulator of tumor dissemination. Future prospective studies with larger patient groups are warranted to validate its clinical utility.

## Conclusions

This study comprehensively investigates the heterogeneity of breast cancer CTCs, the EMT process, and the clinical relevance of MMP9 expression from multiple perspectives. Our findings demonstrate a high prevalence of EMT among CTCs, with a predominance of mixed and mesenchymal phenotypes. The abundance of total CTCs and specific subtypes was closely associated with key clinicopathological parameters: increased CTC loads were linked to advanced clinical stage, larger tumor size, greater lymph node metastasis, and elevated Ki-67 expression, while mesenchymal CTC counts rose specifically with escalating lymph node involvement. Furthermore, MMP9 expression exhibited a dual role, showing a positive correlation with lymph node metastasis yet an inverse relationship with tumor diameter. Critically, we identified a significant interplay between MMP9 and CTC dynamics. Mixed CTCs showed the highest MMP9 positivity rate, and elevated MMP9 expression in tumor tissue promoted CTC generation, EMT progression, and an increase in MMP9-positive CTCs. Collectively, these findings provide a new theoretical basis for precision diagnosis and treatment of breast cancer by elucidating the interconnected roles of CTC heterogeneity and MMP9 in tumor progression.

## Acknowledgements

This study is funded by the National Natural Science Foundation of China (82260423), the Guangxi Education Department (2022KY0494, 2022KY0503). We would like to thank Dr. Changsheng Ye and Dr. Lujia Chen for their guidance and advice for this work.

## Preprints

An early draft of this article has been uploaded in preprints server bioRxiv 2025.06.30.662312; doi: https://doi.org/10.1101/2025.06.30.662312. This article has not been published elsewhere.

## Use of Large Language Models, AI, and Machine Learning Tools

Not applicable.

## Data availability

All data supporting the findings of this study are available from the corresponding author upon reasonable request.

## Template for Ethical and Legal Declarations

Template for Ethical and Legal Declarations

