## Supplemental fig table for "Circulating tumor cell heterogeneity and MMP9 expression in clinical correlation with breast cancer"

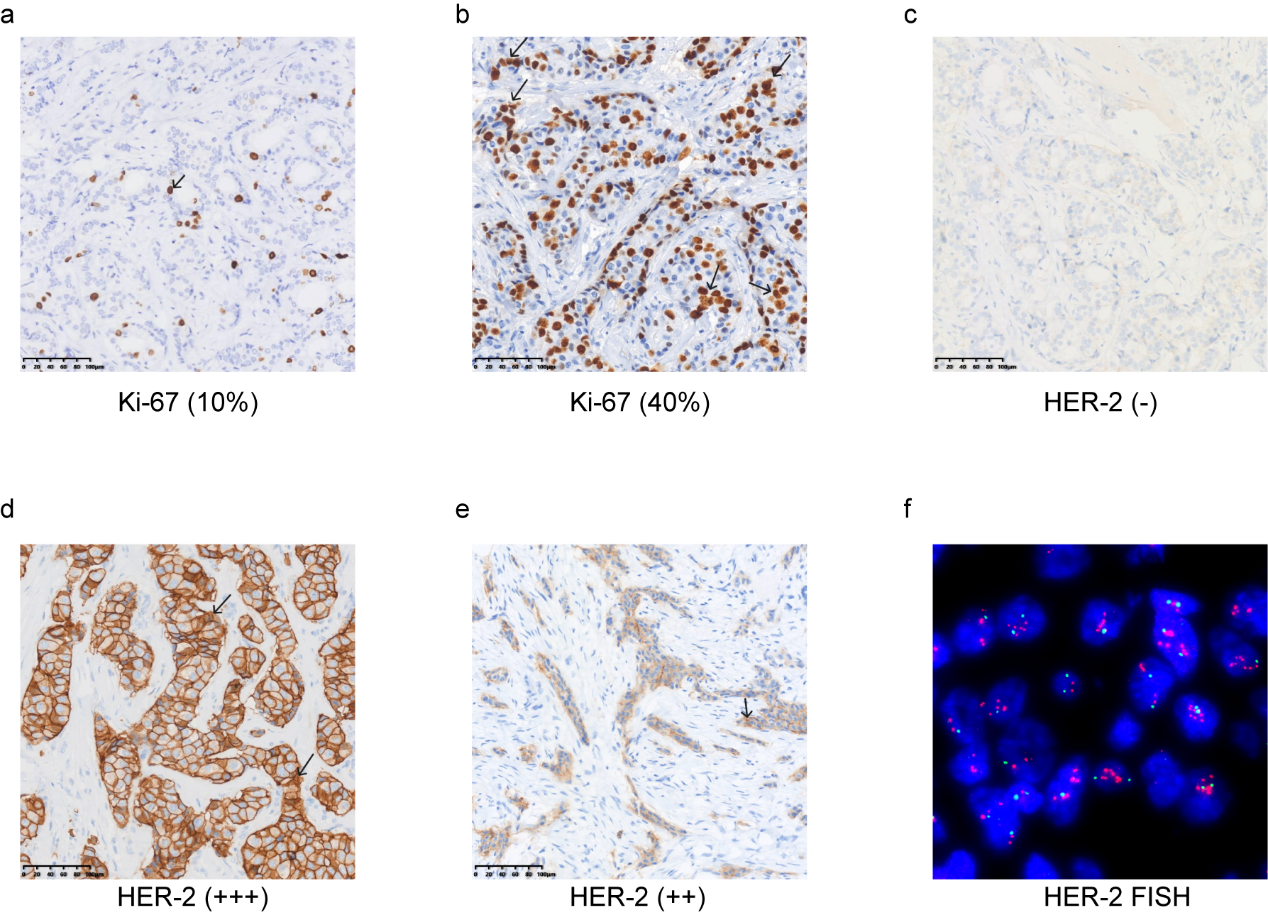

Supplementary figure 1 Representative IHC images of Ki-67+/- and HER-2 +/- tumor tissue from breast cancer patients.

a, b: representative IHC images of Ki-67- (a) and Ki-67+ (b), Ki-67 >20% was deemed positive, arrow points to positive IHC staining cells; c, d, e: representative IHC images of HER-2- (c), HER-2+++ (d) and HER-2++ (e); f: representative FISH image of HER-2, red dots: HER-2 probe; green dots: CEP17 probe. a-e: arrow points to positive IHC staining cells, representative microphotographs at a 200×magnification.

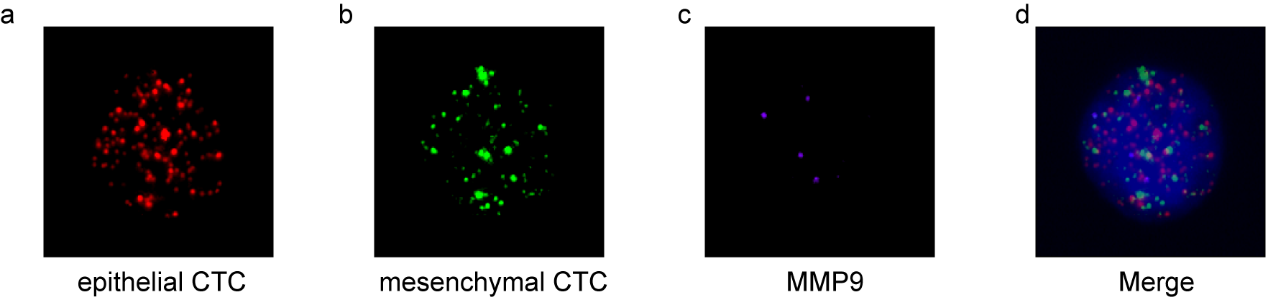

Supplementary figure 2 Representative images of MMP9+ CTCs

a: epithelial CTC (red fluorescence signal); b: mesenchymal CTC (green fluorescence signal); c: MMP9 positive CTCs (purple fluorescence signal); d: merged image.

Supplementary Table 1 Capture probe sequences for CanPatrol^TM^

| Gene | Sequence (5’ - 3’) |
| --- | --- |
| EpCAM | TGGTGCTCGTTGATGAGTCA |
|  | AGCCAGCTTTGAGCAAATGA |
|  | AAAGCCCATCATTGTTCTGG |
|  | CTCTCATCGCAGTCAGGATC |
|  | TCCTTGTCTGTTCTTCTGAC |
|  | CTCAGAGCAGGTTATTTCAG |
| CK8 | CGTACCTTGTCTATGAAGGA |
|  | ACTTGGTCTCCAGCATCTTG |
|  | CCTAAGGTTGTTGATGTAGC |
|  | CTGAGGAAGTTGATCTCGTC |
|  | CAGATGTGTCCGAGATCTGG |
|  | TGACCTCAGCAATGATGCTG |
| CK18 | AGAAAGGACAGGACTCAGGC |
|  | GAGTGGTGAAGCTCATGCTG |
|  | TCAGGTCCTCGATGATCTTG |
|  | CAATCTGCAGAACGATGCGG |
|  | AAGTCATCAGCAGCAAGACG |
|  | CTGCAGTCGTGTGATATTGG |
| CK19 | CTGTAGGAAGTCATGGCGAG |
|  | AAGTCATCTGCAGCCAGACG |
|  | CTGTTCCGTCTCAAACTTGG |
|  | TTCTTCTTCAGGTAGGCCAG |
|  | CTCAGCGTACTGATTTCCTC |
|  | GTGAACCAGGCTTCAGCATC |
| Vimentin | GAGCGAGAGTGGCAGAGGAC |
|  | CTTTGTCGTTGGTTAGCTGG |
|  | CATATTGCTGACGTACGTCA |
|  | GAGCGCCCCTAAGTTTTTAA |
|  | AAGATTGCAGGGTGTTTTCG |
|  | GGCCAATAGTGTCTTGGTAG |
| Twist | ACAATGACATCTAGGTCTCC |
|  | CTGGTAGAGGAAGTCGATGT |
|  | CAACTGTTCAGACTTCTATC |
|  | CCTCTTGAGAATGCATGCAT |
|  | TTTCAGTGGCTGATTGGCAC |
|  | TTACCATGGGTCCTCAATAA |
| CD45 | TCGCAATTCTTATGCGACTC |
|  | TGTCATGGAGACAGTCATGT |
|  | GTATTTCCAGCTTCAACTTC |
|  | CCATCAATATAGCTGGCATT |
|  | TTGTGCAGCAATGTATTTCC |
|  | TACTTGAACCATCAGGCATC |
